# The spatial and temporal dynamics of the nuclear RNAi-targeted RNA transcripts in *Caenorhabditis elegans*

**DOI:** 10.1101/303230

**Authors:** Julie Zhouli Ni, Natallia Kalinava, Sofia Galindo Mendoza, Sam Guoping Gu

## Abstract

Small RNA-guided chromatin silencing, also referred to as nuclear RNAi, plays an essential role in genome surveillance in eukaryotes and provides a unique paradigm to explore the complexity in RNA-mediated chromatin regulation and transgenerational epigenetics. A well-recognized paradox in this research area is that transcription of the target loci is necessary for the initiation and maintenance of the silencing at the same loci. How the two opposing activities (transcriptional activation and repression) are coordinated during animal development is poorly understood. To resolve this gap, we took single-molecule RNA imaging, deep-sequencing, and genetic approaches towards delineating the developmental regulation and subcellular localization of RNA transcripts of two exemplary endogenous germline nuclear RNAi targets in *C. elegans*, Cer3 and Cer8 LTR retrotransposons. By examining the wild type and a collection of mutant strains, we found that transcription and silencing cycle of Cer3 and Cer8 is tightly coupled with the early embryogenesis and germline mitotic and meiotic cell cycles. Strikingly, Cer3 and Cer8 transcripts are exclusively localized in the nuclei of germ cells in both wild type and germline nuclear RNAi-defective mutant animals. RNA-sequencing analysis found that this nuclear enrichment feature is a general feature for the endogenous targets of the germline nuclear RNAi pathway. In addition, the germline and somatic repressions of Cer3 have different genetic requirement for the three H3K9 histone methyltransferases, MET-2, SET-25, and SET-32, in conjunction with the nuclear Argonaute protein WAGO-9/HRDE-1. These results provide a first comprehensive cellular and developmental characterization of the nuclear RNAi-targeted endogenous targets throughout animal reproductive cycle. Altogether, these results support a model in which (1) both the transcriptional activation and repression steps of the germline nuclear RNAi pathway are tightly coupled with animal development, (2) the endogenous targets exhibit a hallmark of nuclear enrichment of their transcripts, and (3) different heterochromatin enzymes play distinct roles in somatic and germline silencing of the endogenous targets.

## Introduction

Besides the classical RNAi pathway, in which a target gene is silenced by mRNA degradation, small RNA can also guide heterochromatin formation and transcriptional silencing through the nuclear RNAi pathway in a diverse set of eukaryotic organisms (see [1, 2]for reviews). In *Caenorhabditis elegans*, nuclear RNAi can be conveniently triggered by exogenous dsRNA and result in highly specific heterochromatic responses (H3K9me3 and H3K27me3) at a target gene[3, 4]. Interestingly, both the transcriptional repression and heterochromatin marks at a target gene can persist for multiple generations after the dsRNA exposure has been removed [3, 4]. These features, together with a ~3-day reproductive cycle (23-25°C) and the powerful genetics, make *C. elegans* a highly tractable system to study RNA-mediated chromatin regulation and transgenerational epigenetics in animals.

In order to establish a framework to study the native functions and mechanisms of nuclear RNAi in *C. elegans*, we previously identified the endogenous targets of the germline nuclear RNAi pathway. These endogenous targets were defined by the presence of abundant 22G-class siRNAs, and the H3K9me3 or the transcription repressive state that is dependent on the germline nuclear AGO protein HRDE-1(<u>h</u>eritable <u>R</u>NAi <u>de</u>fective 1). The most prominent endogenous targets include LTR retrotransposons in the *C. elegans*’s genome [5, 6]. These LTR retrotransposons have strong transcription potentials, evidenced by their high RNA polymerase II (Pol II) occupancy and RNA transcripts levels in the germline nuclear RNA-defective mutants[5, 6]. In wild type animals, the transcription activities of these LTR retrotransposons are repressed by nuclear RNAi to an extremely low level[5, 6]. Despite the low expression, transcription of these endogenous targets is expected to play an essential role for their silencing because (1) the RNA transcripts are needed as templates for the siRNA biogenesis [7, 8], and (2) the nascent transcripts also function as the scaffold for the binding of siRNA-guided silencing complexes[9, 10]. These features form an intriguing paradox: the silencing of nuclear RNAi targets is dependent on the expression of the same loci. In plants and *S. pombe*, this paradox is solved by coupling the expression of the nuclear RNAi target loci to distinct cell type or cell cycle stages [11–13]. Some small RNA targeted and epigenetically silenced transposable elements (TEs) transiently expressed at specific developmental stages and in specific cells during normal development of wild type plants and animals[14–16], a phenomenon referred to as developmental relaxation of TE silencing (DRTS) [17]. DRTS is critical to maintain TE silencing by providing siRNAs to reinforce TE silencing [13, 18, 19], in addition, it also benefit the host by providing innate immunity against similar TEs or viruses [20, 21].

When and where the endogenous targets of nuclear RNAi are expressed in the wild type *C. elegans*, as well as nuclear RNAi-defective mutant strains, is virtually unknown at the cellular level. In addition, although many *nrde* and *hrde* genes (named after the nuclear RNAi- and heritable RNAi-defective phenotypes) have been identified in recent years, the dynamics of this pathway at the subcellular level, as well as in the context of animal development is not well understood. We are particularly interested in determining how the two seemly conflicting activities, HRDE-1-dependent transcriptional repression and expression of the target transcripts are coordinated during the reproductive cycle. *C. elegans* is well suited for this because the transparent body, a well-organized developmental pattern, and a large collection of mutant strains.

We and others recently determined that the H3K9me3 at the HRDE-1 targets requires three histone methyltransferases (HMTs), MET-2, SET-25 and SET-32[4, 22–24]. Combined genetic and genomic analyses revealed a highly complex relationship between H3K9 HMTs and HRDE-1 in their roles in nuclear RNAi. Mutant animals that lack H3K9me3 due to combined mutations in all three H3K9 HMT genes do not exhibit any activation of the endogenous targets of HRDE-1, indicating that HRDE-1-dependent transcriptional repression can occur in a H3K9me3-independent manner. On the other hand, mutant animals with combined loss of H3K9 HMTs and HRDE-1 have much higher transcription activities of the endogenous targets than the *hrde-1* mutant animals (Natallia Kalinava, unpublished data). This indicates that H3K9me3 does play a repressive role at the endogenous targets, at least when certain HRDE-1-dependent functions are compromised. However, when and where the H3K9me3-dependent repression occurs during animal development is unknown.

By taking the single-molecule fluorescent in situ hybridization (smFISH) approach, here we delineated the developmental expression of the Cer3 and Cer8 RNA transcripts at the single-cell resolution in wild type and a variety of mutant strains. We found that both the transcriptional relaxation (observed in the wild type) and repression (inferred by using various silencing defective mutants) of the endogenous targets are tightly linked to animal development. Somatic and germline silencing have distinct requirement for the H3K9 HMTs. In addition, we identified nuclear enrichment as a hallmark for nuclear RNAi-targeted endogenous transcripts.

## Result

### Background

*C. elegans* germline development begins with the birth of the founder primordial germ cell (PGC) after the initial four embryonic cell divisions[25]. The founder PGC divides only once to produces two daughter PGCs during embryogenesis. Starting from the end of the first larval (L1) stage, PGCs undergo continuous proliferation to produce ~ 1000 germ cells in each of the two gonads at the adult stage. When a hermaphrodite animal reaches the L3 stage, germ cells at the proximal region of the gonad enter meiosis and spermatogenesis, while germ cells at the distal region of the gonad continue to divide mitotically. Such polarity of the germline was maintained in L4 and adult hermaphrodite animals, except that the proximal germ cells switch to oogenesis.

An adult hermaphrodite has two symmetric U-shaped gonad arms. Different developmental stages of germ cells - proliferation, transition zone, pachytene, diplotene, and diakinesis – are temporally and spatially organized along the distal-to-proximal axis of each gonad arm, and can be distinguished by their locations in the gonad, cellular morphology, as well as their chromosome morphology (Fig. 1A). These features, together with the powerful genetics and a transparent body of *C. elegans*, provide a highly tractable system to study germline nuclear RNAi at the single-cell resolution and in the context of animal development.

**Figure 1.**
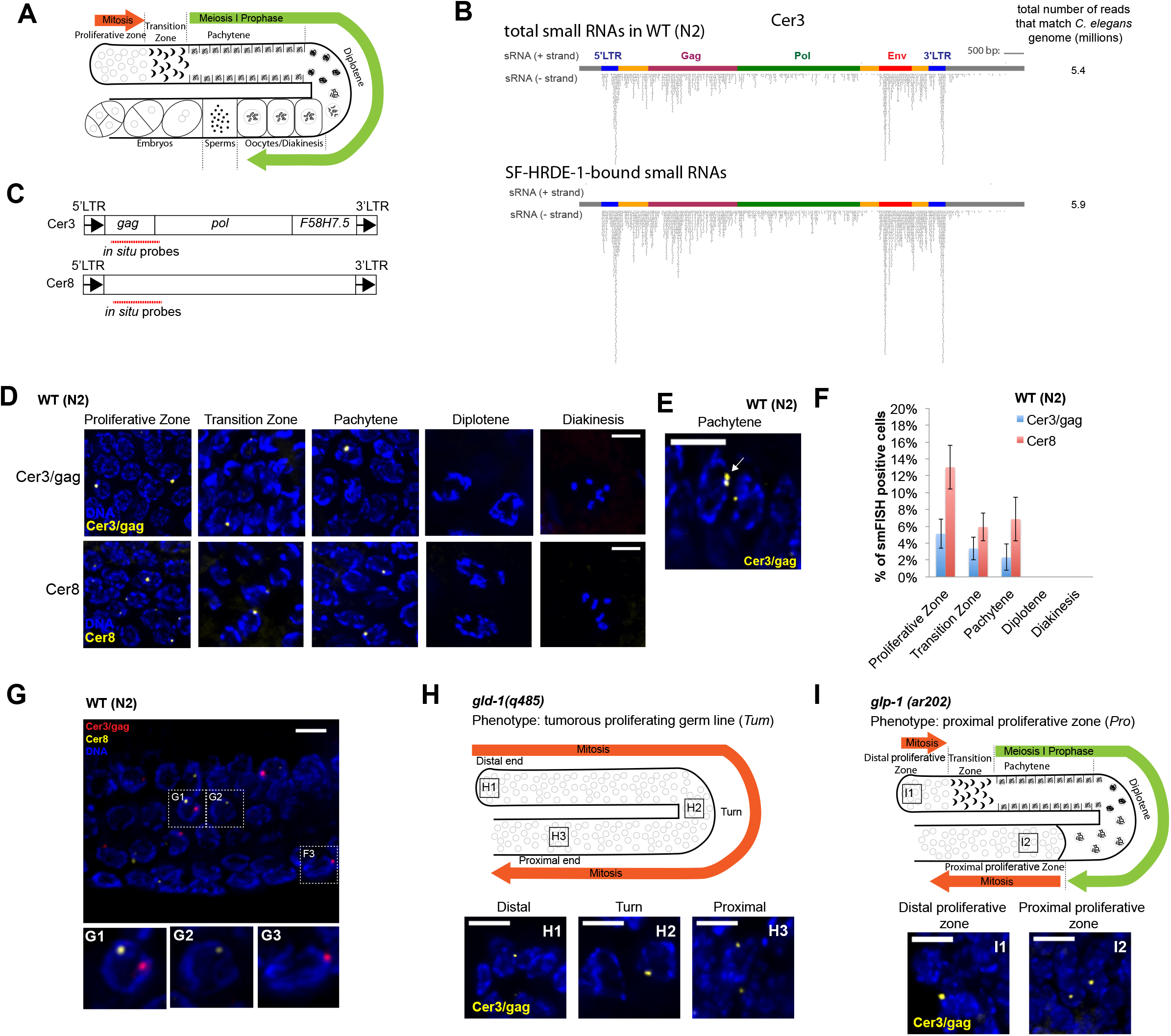
smFISH analysis of Cer3 and Cer8 RNA transcripts in wild type adult germline. (**A**) A schematic diagram of the gonad in *C. elegans* hermaphrodite adult. (**B**) Total siRNAs and HRDE-1-co-IP siRNAs at Cer3. Each dot represents a sequenced siRNA. Sense and antisense siRNAs were plotted separately as indicated. (**C**) Design of Cer3/gag and Cer8 smFISH probe sets. (**D**) Cer3/gag and Cer8 smFISH signals at different stages of adult germ cells. Wild type (N2) hermaphrodite animals were used (also for panels **E-G**). (**E**) An example of a pair of adjacent Cer3/gag smFISH dots (white arrow) in pachytene nucleus. (**F**) Percentage of Cer3/gag or Cer8 smFISH positive nuclei observed at different developmental stages of adult germ cells. (**G**) Two-color smFISH analysis of Cer3/gag and Cer8 in the pachytene stage. (**H**) Schematic diagram of *gld-1(q485)* tumorous germline phenotype and Cer3/gag smFISH at different regions of *gld-1(q485)* gonad. (**I**) Schematic diagram of *glp-1(ar202)* proximal proliferative zone and Cer3/gag smFISH at different regions of *glp-1(ar202)* gonad. See Fig. S4 for smFISH profiles of the entire gonads. (Scale bars are 5 μm.)

### Cer3 LTR retrotransposon as a model system to study nuclear RNAi

The *gypsy-family* LTR retrotransposon Cer3 was chosen as the primary target in this study for the following reasons: (1) Cer3 is a prominent endogenous target of the germline nuclear RNAi pathway. It is transcriptionally repressed in a HRDE-1-dependent manner and exhibits a high level of H3K9me3[5]. Cer3 is also associated with abundant endo-siRNAs, which are bound by HRDE-1 (Fig. 1B and S1). (2) There is only one allele of Cer3 in the haploid genome of the N2 laboratory strain (ChrIV:912948-921667, WS190)[26, 27], which makes it highly tractable for molecular and imaging analyses. (3) The 5’ and 3’ LTRs (424 bases) are identical, suggesting that its entry into the *C. elegans* genome is sufficiently recent and the original *cis* regulatory elements are likely to be intact.

### Cer3 is transcribed in a small subset of germline nuclei in wild type adults

To characterize the expression pattern of Cer3 RNA transcripts at the single-cell resolution, we performed smFISH [28, 29] against the Cer3 *gag* gene (Fig. 1C). To evaluate the experimental design, we initially used *hrde-1* mutant animals and a Cer3-deleted wild isolate (JU533) [30] for the smFISH analysis. We detected robust signals in the germline, but not the somatic cells, of *hrde-1* mutant adults (Figure 2A and Fig S2, we will provide a more detailed characterization in a later section) and no signal in the JU533 strain (Fig S2). Therefore, our Cer3 smFISH signals are specific to the Cer3 RNA transcripts. We will refer to cells that are positive for the Cer3 smFISH signals as Cer3+ cells in the rest of the paper.

**Figure 2.**
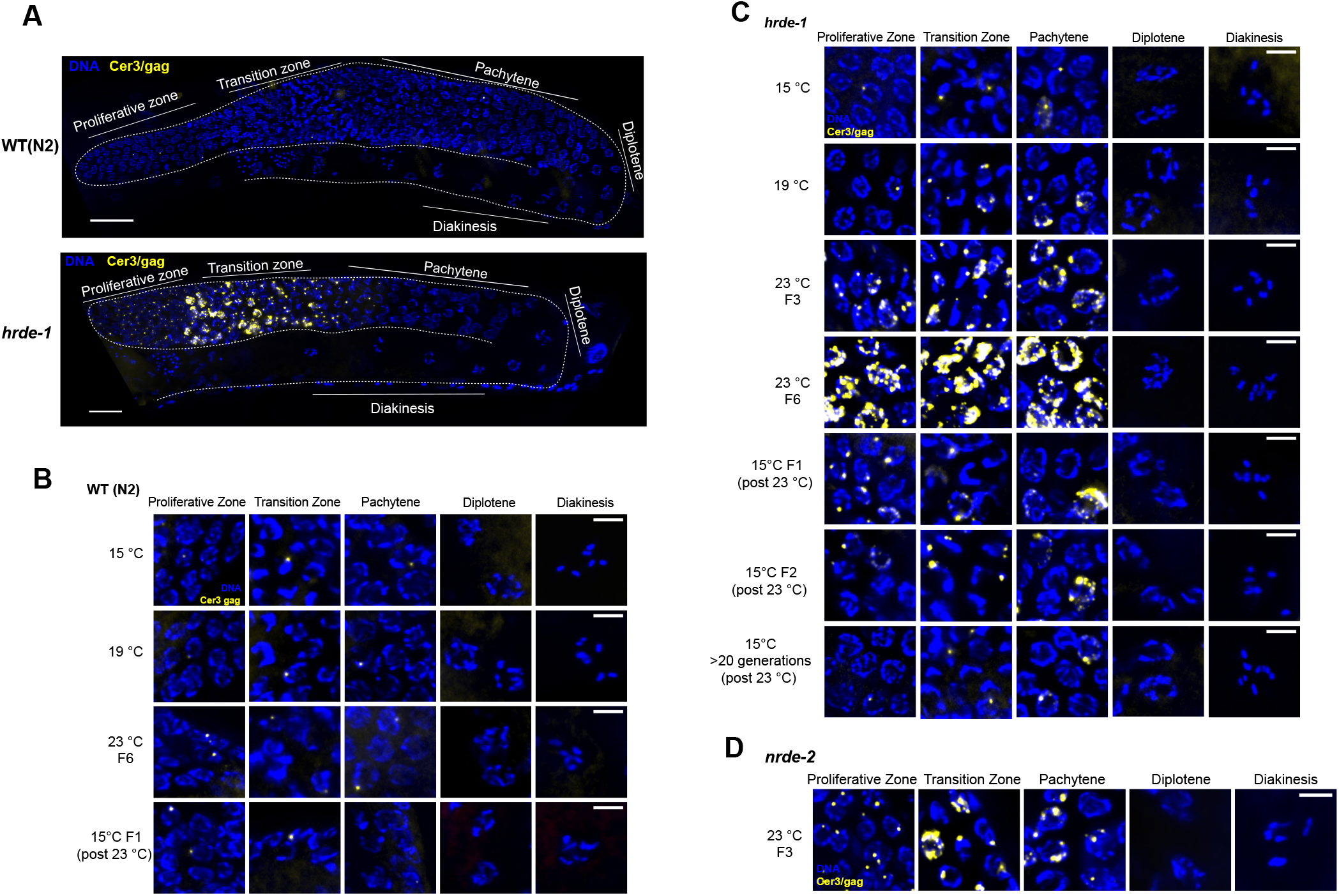
Cer3 smFISH analysis in germline nuclear RNAi defective mutants and the effect of temperature on Cer3 expression. (**A**) Cer3/gag smFISH in wild-type (N2) and *hrde-1* mutant adult. Gonads are outlined. Scale bars are 20 μm. (**B**) Cer3/gag smFISH at different germline developmental stages of wild type (N2) animals that were cultured at 15°C, 19°C, 23°C, or 15°C F1 (the first generation after culturing at 23°C for six generations). Scale bars are 5 μm. (**C**) Cer3/gag smFISH at different germline developmental stages of *hrde-1* adults that were cultured at 15°C, 19°C, 23°C F3/F6 (F3 and F6 at 23°C after shifting from >20 generations growth at 15°C), and 15°C post 23°C (F1, F2, and >20 generations at 15°C after shifting from 6 generations of growth at 23°C). Scale bars are 5μm. (**D**) Cer3 gag smFISH for *nrde-2* mutant adult animals cultured at 23°C for three generations. Scale bars are 5μm.

In wild type adult hermaphrodite animals (23°C), the Cer3 smFISH signals were restricted to a small fraction of germ cell nuclei and absent in any somatic cells (n=10 animals). The distribution of the Cer3+ cells in the adult germline is not uniform. They were present in approximately 5.1%, 3.4%, and 2.3% of the nuclei in the proliferative zone, transition zone, and pachytene zone, and were absent in the post-pachytene germ nuclei, including diplotene and oocytes (n=5 gonads) (Fig. 1C-D). Such polarization was also present in the male germline (data not shown). In both male and female germline, the Cer3 smFISH signal was absent in the shared cytoplasm core of the gonad.

There was usually just one smFISH dot in a Cer3+ nucleus, and occasionally a pair of adjacent dots in the pachytene nuclei (Fig. 1E). We never observed nuclei with three or more dots. These observations suggest that the Cer3 smFISH signals in wild type animals correspond to the nascent Cer3 RNAs at the transcription sites.

### Co-smFISH analysis of two different endogenous targets of germline nuclear RNAi

We also examined a second endogenous target of the germline nuclear RNAi pathway, the Pao-family LTR retrotransposon Cer8. The Cer8 smFISH signals were highly similar to Cer3 in the overall morphology and abundance of smFISH dots (one or two per Cer8+ cell), and the tissue-specificity (Fig. 1C and 1D). Cer8+ cells were approximately 2-3 fold more abundant than the number of Cer3+ cells (Fig. 1F).

We then asked how often Cer3 and Cer8 are expressed in the same cells. To this end, we performed two-color smFISH against Cer3 and Cer8 transcripts in wild type adult animals (Fig. 1G). We found that Cer3+Cer8+ germ cells were rare (1.2%, 0.5%, 0.9% of the germ nuclei in proliferative zone, transition zone, and pachytene, receptively, Fisher exact test: P>0.05). In the few double positive cells, Cer3 and Cer8 smFISH dots did not co-localize(Fig. 1G).

Based on these results, we propose the following model. Cer3 and Cer8 LTR retrotransposons are not fully silenced at the transcriptional level in wild type animals. Transcriptional relaxation of Cer3 and Cer8 occurs in at least a subset of proliferating germ cells and early meiotic cells. Cer3 and Cer8 RNA transcripts do not accumulate beyond the transcription site. In addition, the transcripts do not persist in the late meiotic cells, suggesting that Cer3 and Cer8 are subject to nuclear degradation once they are transcribed. Cer3 or Cer8 are completely silenced in the adult somatic cells. Consistent with this germline-specific expression profile, the Cer3 and Cer8 siRNAs, as well as siRNAs of other endogenous targets of HRDE-1, are enriched in the germline by using sRNA-seq data in Gent, et al., [31] (Fig. S2).

### Cer3 expression is linked to mitotic proliferation

To further explore whether the transcriptional relaxation of Cer3 is linked to mitotic proliferation, we performed Cer3/gag smFISH in the *gld-1(q485)* mutant strain, in which germ cells are defective in the mitotic-to-meiotic transition and all germ cells undergo mitotic cycling. This results in a tumorous germline that contains predominantly the mitotic nuclei throughout the gonad [32, 33]. We found that the small number of Cer3+ nuclei were distributed throughout the entire length of the *gld-1* tumorous gonad (Fig. 1H and S4). We also examined a second germline mutant strain, *glp-1(ar202)*, in which proximal germ cells exit meiotic differentiation and re-enter mitotic cycling at the restrictive temperature (25°C)[34]. We found that Cer3+ nuclei in this mutant were again correlated with the proliferative cells: they were present in the proximal end of gonad arms where meiotic prophase germ cells switched back to the mitotic fate (Fig. 1I and S4), in addition to the distal germ nuclei. We did not observe any Cer3+ nuclei at the diplotene stage in *glp-1(ar202)*. These results, together with aforementioned results of WT animals, indicate that Cer3 expression is developmentally regulated in the germline: Cer3 transcription is partially activated in mitotic proliferating cells and cells in the early stages of meiotic prophase I till pachytene, but completely repressed in the post-pachytene meiotic cells.

### HRDE-1-dependent transcriptional silencing occurs at specific germline developmental stages

HRDE-1 protein is localized in all germ cell nuclei in a hermaphrodite adult, except sperms[23, 35, 36] (Fig. S1). However, it was unknown when and where HRDE-1-mediated silencing occurs. Does HRDE-1-dependent silencing occur in all HRDE-1-expressing germ cells? Does HRDE-1-dependent silencing occur in non-HRDE-1-expressing cells? To answer these questions, we performed Cer3 smFISH analysis in *hrde-1* mutant adults (23°C). Consistent with our previous RNA-seq analysis[5], the Cer3 smFISH signals were dramatically increased in *hrde-1* mutants compared to WT (Fig. 2A). Strikingly, the smFISH signals in *hrde-1* mutant adult gonads were exclusively localized in the nuclei (Fig. 2C and 3A). No Cer3 smFISH signals were detected in the shared cytoplasmic core. As a control experiment, we detected abundant mRNA of the germline-specific gene *oma-1* in the cytoplasmic core by smFISH (Fig. 3A).

**Figure 3.**
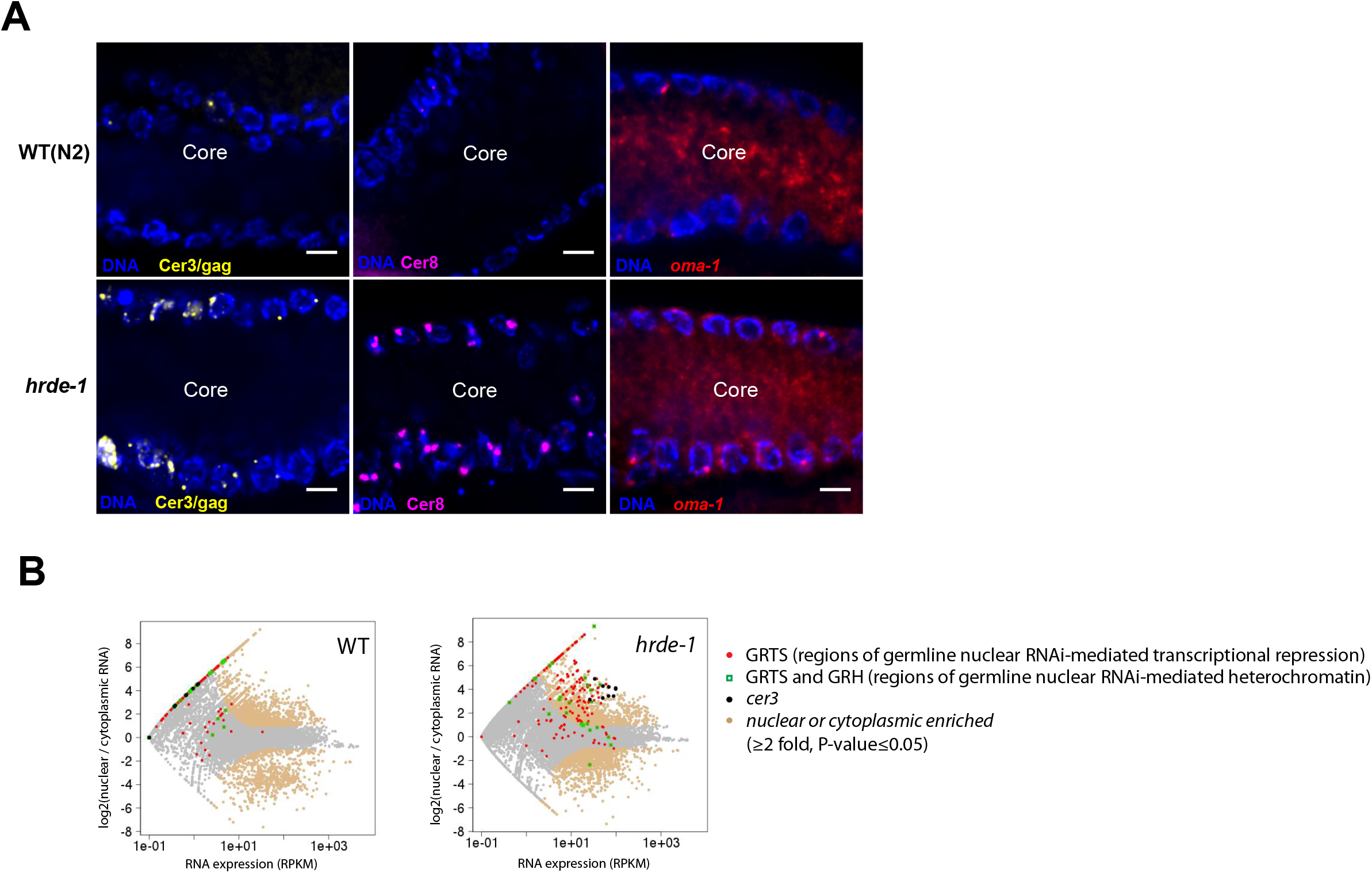
Transcripts of endogenous nuclear RNAi targets are nuclear enriched. (**A**) smFISH against Cer3/gag, Cer8, and *oma-1* in adult wild type (N2) and *hrde-1* pachytene germline. The image shows a representative longitudinal section through the gonad core. Note that Cer3/gag and Cer8 smFISH were only present in the nuclei, and were absent in the cytoplasm core. (**B**) RNA-seq analysis of nuclear and cytoplasmic RNA samples from WT (N2) and *hrde-1* mutant adults. The average nuclear and cytoplasmic RNA expression levels (RPKM, x-axis) and the log2 ratios of nuclear to cytoplasmic RNA level (y-axis) are plotted for all 1-kb regions in the genome. Regions with at least 2-fold difference between nuclear and cytoplasm RNA (p-value⩽0.05) are colored in beige.

Most of the distal germ nuclei (progenitor zone, transition zone, and early pachytene nuclei) in *hrde-1* mutants exhibited the Cer3 smFISH signals (Fig. 2A and 2C). In addition, the average smFISH signal in Cer3+ nuclei in *hrde-1* mutant was much stronger than WT. The strongest signals appeared in the transition zone, where larger dots or irregular patches of Cer3/gag signals were frequently observed.

Despite the strong Cer3 RNA expression in the distal regions of the gonad, Cer3 smFISH signals were absent in the late pachytene, diplotene, and diakinesis stages in *hrde-1* mutants. No Cer3 RNA transcript was detected in *hrde-1* adult somatic cells. The same profile was observed for Cer8 as well (Fig. S5). We also examined Cer3 in a second germline nuclear RNAi defective strain that has the *nrde-2* loss-of-function mutation[9]. We previously found that the endogenous HRDE-1 targets are also derepressed in the *nrde-2* mutants[5]. Here we found that the *nrde-2* mutant strain exhibited the same Cer3 smFISH profile in the gonad as *hrde-1* mutant (Fig. 2D). These results indicate that the germline nuclear RNAi pathway is essential for repression of Cer3 and Cer8 in the distal germline regions (progenitor zone, transition zone, and early pachytene) in adult germline tissue, but is dispensable in the proximal germline regions (late pachytene, diplotene and diakinesis). A germline nuclear RNAi-independent mechanism is likely the cause of the transcriptional repressive states of Cer3 and Cer8 in the proximal germline regions.

### Heat stress-induced transgenerational accumulation of Cer3 transcripts

We previously found that heat stress enhances the transcription of Cer3 and many other endogenous targets in *hrde-1* mutant animals[6]. To determine the cellular origin of the heat-induced activation, we performed Cer3 smFISH using *hrde-1* mutants that had been cultured at different temperatures: two permissive temperatures at 15°C and 19°C and a restrictive temperature at 23°C. To capture the transgenerational effect caused by heat stress, we used *hrde-1* mutants that had been continuously cultured at 23°C for three and six generations (referred to as 23°C F3 and 23°C F6 samples). (The ancestral population was cultured at 15°C.)

Consistent with our previously RNA-seq analysis, *hrde-1* mutant animals exhibited much higher Cer3 smFISH signals at 23°C than 15°C and 19°C (Fig. 2C). We did not detect any heat-induced enhancement of Cer3 expression in wild-type animals using smFISH (Fig. 2B). The 23°C F6 *hrde-1* mutant animals also showed much stronger smFISH signals than the 23°C F3 mutant adults. Such transgenerationally progressive effect was also observed by our previous RNA-seq analysis[6]. Our smFISH results revealed that the enhanced activation of Cer3 was primarily due to higher expressions in the individual Cer3+ nuclei, rather than more Cer3+ nuclei, in the 23°C F6 samples compared to the other samples. The Cer3+ nuclei in *hrde-1* mutant strain were still limited to the distal gonad in the 23°C F6 samples (Fig. 2C). These results, together with our previous ChIP-seq analysis showing increased Pol II occupancy at Cer3 by heat stress, indicate that Cer3 transcription is enhanced by heat in *hrde-1* mutants, and the temperature effect is transgenerationally progressive.

A significant number of adult germ cells in wild type animals undergo apoptosis[37]. To investigate whether the germline degeneration of *hrde-1* mutants is caused by increased frequency of apoptosis, we examined germ cell corpses using the CED-1:GFP reporter[38]. We also performed Cer3 smFISH in the same animals to test the possible link between Cer3 derepression and apoptosis. We found that the frequency and distribution of CED-1:GFP-labeled cells in *hrde-1* mutants were similar to wild-type(Fig. S6). In addition, CED-1:GFP-labeled cells in both wild type and *hrde-1* mutant gonads did not have any Cer3 smFISH signals. Therefore, *hrde-1* mutant is not associated with enhanced apoptosis of meiotic germ cells. Our results also argue against the role of Cer3 derepression in promoting the apoptosis of germ cells.

### Transcripts of the endogenous HRDE-1 targets are enriched in the nuclei of adult germline

Our smFISH analysis found that Cer3 and Cer8 RNA transcripts were only present in the nuclei, not in the cytoplasm, in both wild type or *hrde-1* mutant adult germ cells (Fig. 3A). To determine whether the nuclear localization is a general feature for the germline nuclear RNAi-targeted transcripts, we performed RNA-seq for isolated nuclei, cytoplasmic extract, and whole animals of wild type and *hrde-1* mutant adults (23°C for both). To capture both polyadenylated and non-polyadenylated RNA, RNA was sequenced without using the oligo(dT)-based enrichment (referred to as RNA-seq in this work). The nuclear enrichment index (RNA_nuclear_/RNA_cytoplasmic_) was calculated for each 1kb region throughout the genome. By using a two-fold cutoff (p≤0.05), we found that transcripts from 4909 kb and 3669 kb regions were enriched in the nucleus in WT and *hrde-1* mutant, respectively. Introns, particularly the long ones (>2kb), were highly enriched in the nuclear RNA compared to the cytoplasmic RNA, confirming our fractionation procedure (Fig. S7).

We previously classified the endogenous HRDE-1 targets based on the effect of the *hrde-1* mutation at these loci: regions termed germline nuclear RNAi-dependent transcriptional repression (or GRTS) become transcriptionally derepressed in the *hrde-1* mutant, and regions termed germline nuclear RNAi-dependent heterochromatin (or GRH) lose H3K9me3 in the *hrde-1* mutant[5]. We found that many GRTS transcripts were enriched in the nucleus over cytoplasm for both WT and *hrde-1* mutant samples (Fig. 3B). This feature was much more prominent in *hrde-1* mutant than WT animals. In *hrde-1* mutants, 94.8% of the GRTS transcripts were enriched in the nucleus (cutoff: two-fold enrichment and P-values≤0.05). The average fold of nuclear enrichment for the GRTS regions in *hrde-1* mutant was 11.0 (P-value<2.2×10^−16^, Wilcoxon rank-sum test). In WT, 27.7% of the GRTS regions showed the nuclear enrichment using the same cutoff. The lower percentage is likely due to the lower expression of the GRTS loci in WT, which results in higher P-values. Nevertheless, GRTS regions in WT were generally enriched in the nucleus over the cytoplasm with an average nuclear enrichment of 8.1 (P-value<2.2×10^−16^, Wilcoxon rank-sum test).

We found that many GRH transcripts were also enriched in the isolated nuclei of WT and *hrde-1* mutant (Fig. S6). The average nuclear enrichment levels for these regions were 8.9 and 4.5 (both P-values<2.2×10^−16^, Wilcoxon rank-sum test). However, GRH transcripts had a lower degree of nuclear enrichment than GRTS transcripts. 20.9% and 34.0% of GRH regions in WT and *hrde-1* mutant showed at least 2x nuclear enrichment (P-values≤0.05).

These results indicate that a large fraction of the endogenous germline nuclear RNAi targets are enriched in the nucleus, particularly the GRTS transcripts. This nuclear localization feature is not dependent on HRDE-1.

### Distinct genetic requirements of Cer3 silencing in germline and soma

In our recent study using RNA-seq and Pol II ChIP-seq, we found that *H3K9 HMT;hrde-1* compound mutants exhibit stronger Cer3 expression than *hrde-1* single mutant. This was unexpected because the mutations in the *H3K9 HMT* genes, either single or in combination, do not cause any silencing defects at the endogenous HRDE-1 targets, including Cer3. To further investigate the synthetic effect at the single-cell resolution, we performed Cer3 smFISH in WT, *hrde-1* single mutant, *set-32;met-2 set-25* triple mutant and *set-32; met-2 set-25 hrde-1* quadruple mutant adult animals (cultured at 23°C for two generations for all strains). Consistent with our sequencing results, Cer3 smFISH signals in *set-32;met-2 set-25* triple mutants were very low, similar to WT. In contrast, *set-32; met-2 set-25;hrde-1* quadruple mutant animals exhibited dramatically enhanced Cer3 smFISH signals compared to the *hrde-1* single mutant. The Cer3 smFISH signals in the quadruple mutant germline were limited to the proliferative zone, transition zone, and pachytene. Interestingly, intestine nuclei exhibited robust Cer3 smFISH signals in the quadruple mutant (Fig. 4B). Cer3 smFISH signal is also observed in a subset of somatic cells near the head, tail, and epidermis of the quadruple mutant (data not shown), but the identity of these somatic cells can not be further defined based on our DAPI stain of nuclear morphology. None of the WT, *hrde-1* single mutant, and *set-32;met-2 set-25* triple mutant had any Cer3 smFISH signals in adult somatic cells (Fig. 4B).

**Figure 4.**
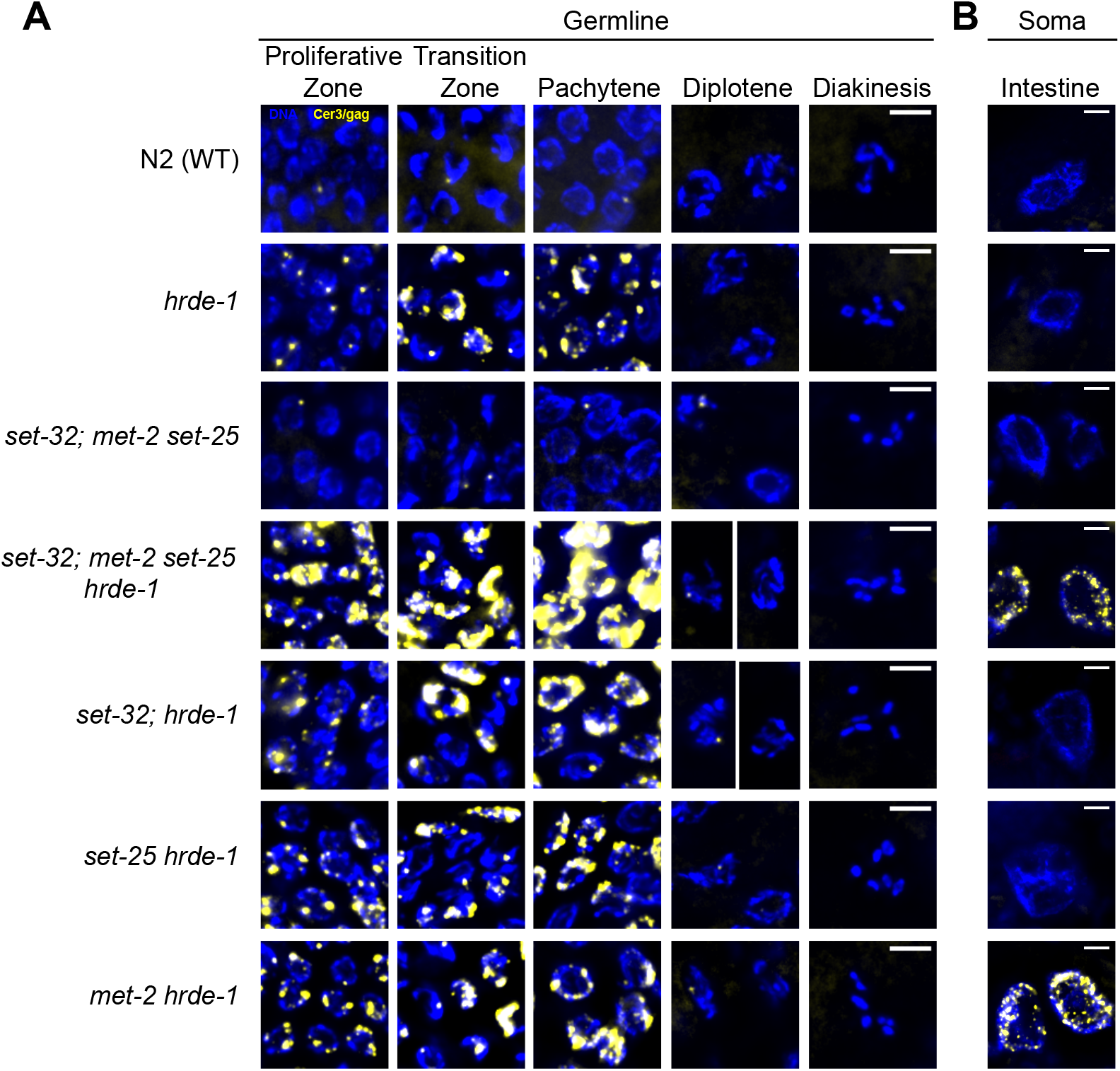
Requirement of different putative H3K9 histone methyltransferases (HMTs) for Cer3 silencing in germline and soma. Cer3/gag smFISH analysis in (**A**) adult germline and (**B**) adult intestinal cells of WT and different mutant strains as indicated. All strains were cultured at 23°C for two generations for this experiment. Scale bars are 5 μm.

To determine which H3K9 HMT contributes to the enhanced Cer3 derepression in germline or soma, we performed Cer3 smFISH in different *H3K9 HMT;hrde-1* double mutants. We found that all three double mutants, *set-32;hrde-1, set-25 hrde-1*, and *met-2 hrde-1*, exhibited enhanced Cer3 expressions in the germline (cultured at 23°C for two generations for all strains) (Fig. 4A). *set-32;hrde-1* double mutants had a higher degree of enhancement than the other two double mutants, but lower than the quadruple mutant animals (Fig. 4A). *met-2 hrde-1* is the only double mutant strain that exhibited Cer3 depression in the intestinal cells (Fig. 4B).

Together these data indicate that H3K9 HMTs are involved in the transcriptional repression of Cer3. However, the underlying mechanisms are likely to be complex for the following reasons. (1) The requirement of H3K9 HMTs in Cer3 silencing is conditioned on the loss of HRDE-1 activity. (2) The tissue specificity of different H3K9 HMTs’ function is not identical. In the intestinal cells, only MET-2, an H3K9me2 HMT, is required to maintain the repressive state of Cer3 in the *hrde-1* mutant. In germline, all three H3K9 HMTs contribute to Cer3 repression in the absence of HRDE-1, with SET-32 contributing more than SET-25 or MET-2.

### Dynamic expression of *Cer3* RNA transcripts during embryogenesis

To investigate Cer3 expression during embryogenesis, we performed Cer3/gag smFISH in the wild type embryos. We did not observe any Cer3 smFISH signals in the fertilized eggs and during the first two rounds of cell division, indicating that Cer3 is transcriptionally inactive during these stages. This quiescent state was followed by a burst of Cer3 RNA expression in most, if not all, cells. The broad expression of Cer3 RNA began at the 8-cell stage and peaked at approximately 30-cell stage, in which the cells are still pluripotent blastomeres. Cer3 RNA expression ceased after the organogenesis/morphogenesis stages (Fig. 5A). Afterwards, Cer3 RNA largely disappeared in all of the embryonic cells except the two primordial germ cells (PGCs) (Fig. 5A and 5B). These results indicate that (1) Cer3 RNA is expressed during embryogenesis, (2) the expression is highly dynamic, and (3) there are two modes of Cer3 RNA expression: a transient, non-cell specific expression in the pluripotent blastomeres and a subsequent PGC-specific expression. Interestingly, Cer3 RNAs in blastomeres were localized in both the cytoplasm and nuclei, while Cer3 RNAs in PGCs were restricted to the nuclei, similarly to the post embryonic germ cells.

**Figure 5.**
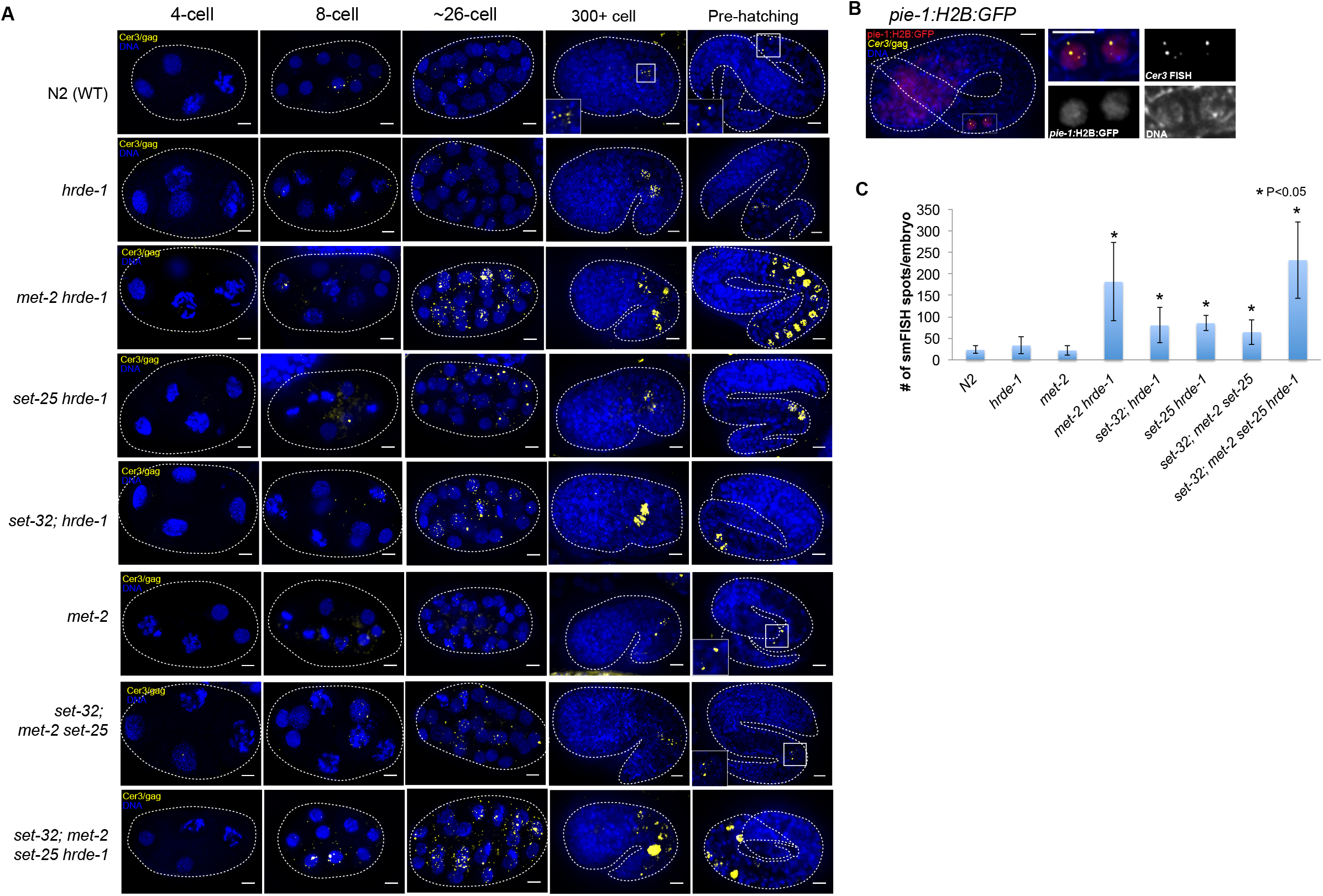
The dynamic and genetic requirement of Cer3 RNA expression during embryogenesis. (**A**) Cer3/gag smFISH in 4-cell, 8-cell, ~26-cell, 300+ cell, and pre-hatching stages of wild-type and different mutant embryos as indicated. (**B**) Simultaneous detection of pie-1 promoter driven GFP::H2B (immunofluorescence with anti-GFP) and Cer3/gag RNA with smFISH in a pre-hatching embryo. The strain carries the Ppie-1::GFP::H2B, but otherwise is wild type. (**C**) Quantification of Cer3/gag smFISH spots in ~26-cell embryos. Scale bars are 5 μm.

To investigate the role of HRDE-1 in Cer3 regulation during embryogenesis, we performed Cer3 smFISH in *hrde-1* embryos. During early embryonic development before the two PGCs were born, Cer3 expression in *hrde-1* is similar as wild-type (Fig. 5A). Cer3 RNA was not expressed in 1-4-cell stage of *hrde-1* mutant embryos, and then become expressed in both nuclei and cytoplasm in many cells from 8-cell to ~30-cell stage. *hrde-1* mutant did not appear to have a higher Cer3 RNA expression at this stage than the WT. During late embryonic development of *hrde-1* mutant, Cer3 RNA expression disappeared in most of the cells just like WT. Compared to WT, *hrde-1* mutant did have higher Cer3 RNA expression in the two PGCs (Fig. 5A). These results indicate that the repressive role of HRDE-1, at least at Cer3, is limited to the PGCs during embryogenesis. An HRDE-1-independent mechanism(s) is likely to repress Cer3 in other embryonic cells.

We then investigated whether any of the three H3K9 HMTs represses Cer3 expression in the somatic or germline lineage in embryos. Compared to the wild-type embryos, *set-32; met2 set-25* embryos had a modest increase in Cer3 RNA at 8-cell to ~30-cell stage, but did not exhibit Cer3 depression in either the somatic lineages or PGCs in later embryos (Fig. 5A and 5C). This suggests that H3K9me3 plays a role in Cer3 repression in the blastomeres during the early embryogenesis, but losing H3K9me3 alone is not sufficient to activate Cer3 in either the somatic cells or PGCs of late embryos.

We then examined *H3K9 HMT/hrde-1* compound mutants. All three double mutants (*met-2 hrde-1, set-25 hrde-1*, and *set-32; hrde-1*), and the quadruple mutant (*set-32; met2 set-25 hrde-1*) have enhanced Cer3 expression in early embryos (8-cell to ~30 cells) compared to WT, suggesting H3K9 HMTs and HRDE-1 in combination play a repressive role for Cer3 expression at this stage. Similar to the adult germline, mutations in H3K9 HMTs enhanced the desilencing of Cer3 in the *hrde-1* mutant background. (Fig. 5A and 5C).

In late embryos of *met-2 hrde-1* and *set-32; met2 set-25 hrde-1* mutants, Cer3 RNA was strongly expressed in a group of nuclei along the epithelial tube where the intestine lineage and PGCs locate [39] (Fig. 5A). However, *met-2* mutation on its own did not cause Cer3 activation in either early or late embryos (Fig. 5A and 5C). Together with enhanced intestine and germline Cer3 expression in *met-2 hrde-1* adult, this data suggest that MET-2 and HRDE-1 synergistically repress Cer3 in both somatic (intestinal) and germline lineages during embryogenesis.

## Discussion

### HRDE-1-dependent repression occurs in the same set of germ nuclei as transcriptional relaxation

Transcriptional relaxation at the endogenous HRDE-1 targets is essential for the reinforcement of their silencing at each generation. The developmental regulation of the transcriptional relaxation in the wild type germline was completely unknown before this study. Transcriptional relaxation can either constantly occur in all germline developmental stages or transiently occur in one or more specific stages. This study found that the latter case is true for the two prominent HRDE-1 targets, Cer3 and Cer8 LTR retrotransposons. Transcriptional relaxation of Cer3 and Cer8 in wild type adult germline is limited to the proliferative germ cells and the first few steps of meiosis prophase I, including leptotene, zygotene, and early pachytene, and does not occur in the subsequent meiotic stages. By examining the Cer3 and Cer8 RNA transcripts in the *hrde-1* mutant, we determined that HRDE-1-dependent repression occurs in the same set of germ nuclei as transcriptional relaxation. Therefore, the transcription and silencing of endogenous nuclear RNAi targets in *C. elegans* occur at the same developmental stages of germline development (summarized in Fig 6A). This different from plants or the fission yeast, in which transcriptional relaxation and nuclear RNAi are separated in different cell types or different stages of the cell cycle[11–13]. These results improve our understanding of germline nuclear RNAi in a number of ways.

**Figure 6.**
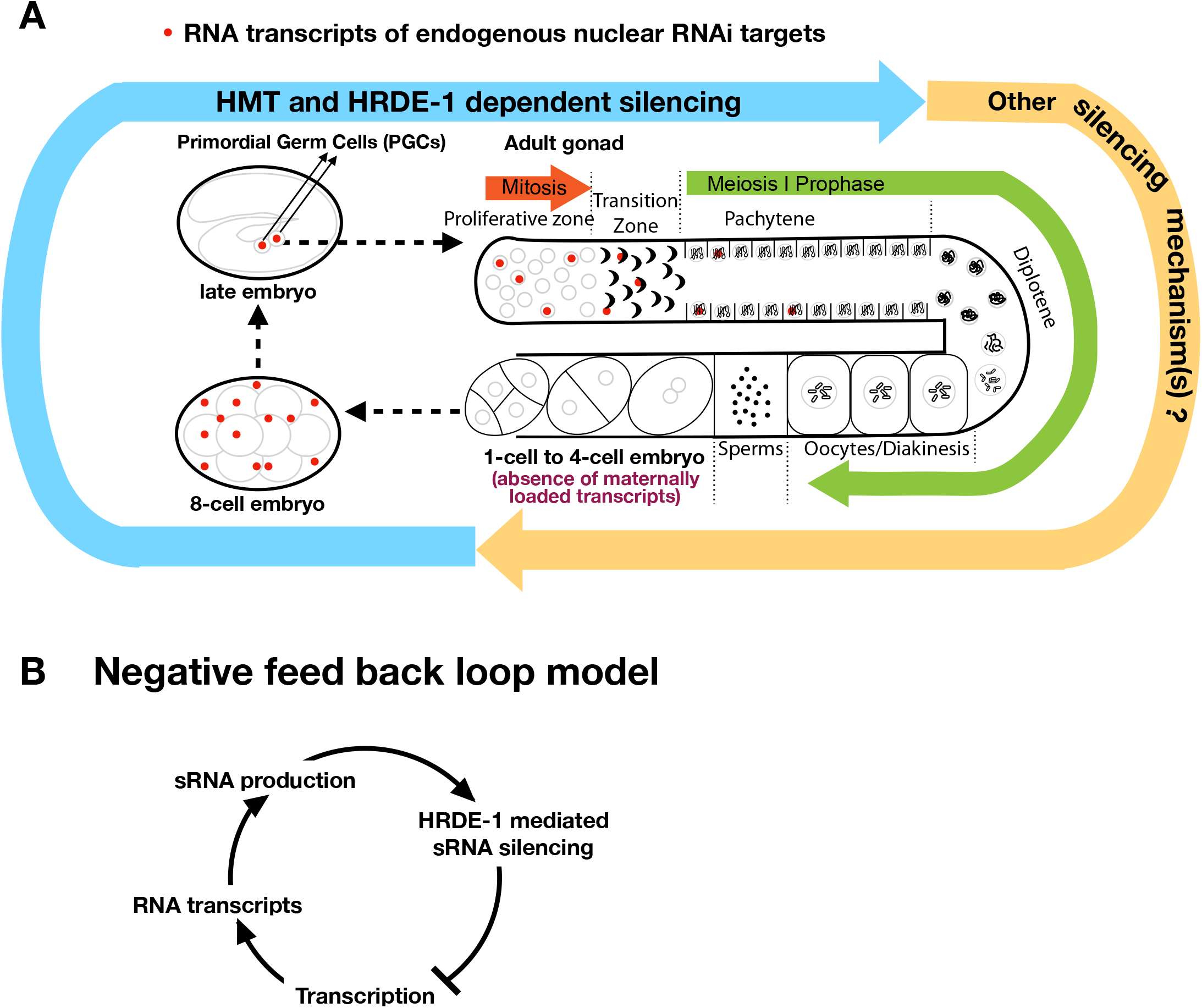
Schematic diagrams of (**A**) Transcription and silencing cycle of endogenous nuclear RNAi targets throughout *C. elegans* reproductive cycle and (**B**) Negative-feed back model providing balance of transcription and silencing of endogenous nuclear RNAi targets at same developmental stage of germ cells.

(1) Because the target RNA transcripts are required as the RNA-dependent RNA polymerase (RdRP) templates for the 22G endo-siRNAs biogenesis, our results suggest that the biogenesis of Cer3 and Cer8 siRNA is limited to the distal regions of the gonad (from proliferative cells to early pachytene). These siRNAs are very likely to persist in the subsequent germline developmental stages and be transmitted to the next generation because of (1) their interactions with HRDE-1 and (2) the presence of HRDE-1 in all female germline developmental stages, including oocyte[23, 35, 36].

(2) Cer3 and Cer8 have strong transcription potentials in distal germ cells. This is evidenced by the Pol II ChIP-seq, RNA-seq[5, 6], and smFISH analyses in *hrde-1* single mutant and *hrde-1 H3K9 HMT* compound mutants. Our previous and this study also showed that heat stress can further enhance the transcription of Cer3 in *hrde-1* mutant, and this effect is transgenerationally progressive[6]. It’ll be of great interest to determine the molecular basis of the transcriptional activations of Cer3 and other endogenous targets.

(3) Transcription relaxation and HRDE-1-mediated repression in principle can form a negative feedback loop (Fig. 6A). Our results suggest that such negative feedback loop occurs in the same cells. Such system can provide a stable silencing of the target loci, even those that have strong transcriptional potentials. For example, an increase in transcription activity will generate more RNA transcripts, which can lead to a higher siRNA production and more targets for HRDE-1-siRNA complex to engage. These changes will in turn lead to a stronger repression. One the other hand, such system avoids a complete silencing (i.e., a complete lack of transcription) because the repressive activity will stop acting on the target gene when there are no transcripts for HRDE-1-siRNA complexes to engage, and then transcription relaxation will occur.

(4) Although HRDE-1 is present in all adult germ nuclei, our analysis indicates that HRDE-1-dependent repression, at least for Cer3 and Cer8, is not needed for the repression in the post-pachytene meiotic cells. This is likely due to the global transcription quiescence state as meiotic cells progress into the late stages, although we cannot rule out another Cer3 or Cer8-specific silencing mechanism beside nuclear RNAi.

### Is nuclear accumulation a sign of “foreign”?

Here we found that, unlike normal mRNAs that are enriched in cytoplasm, transcripts of Cer3, Cer8 and other germline nuclear RNAi targets are enriched in the germline nuclei. This provides a first important cytological hallmark for the endogenous HRDE-1 targets. Nuclear RNA export is coupled with transcription and RNA processing[40]. Aberrant RNAs that fail to be exported are subjected to nuclear RNA degradation. We speculate that nuclear RNA accumulation of the endogenous HRDE-1 targets may reflect aberrant structures or deficiencies in RNA-processing of these transcripts. Such connections have been found in *S. pombe*[41, 42] and recently in *C. elegans*[43, 44]. Investigating the cause of the nuclear enrichment will provide insights into the triggering mechanism of nuclear RNAi in *C. elegans*.

### What is the functional significance of early embryonic transcriptional relaxation?

Our observation of LTR-transcripts in *C. elegans* early embryos has interesting parallel in mammals [20, 45–49]. Retrotransposons are transcribed in human preimplatation embryos at the onset of embryonic genome activation[20, 49], which leads to the formation of viral practical and upregulation of genes in the viral restriction pathway [20]. LTR-derived transcripts are also found in mouse and human stem-cells, which is a major contributor of the stem cell nuclear transcriptome complexity [47, 48]. The actual role of LTR-derived transcription in early *C. elegans* embryo is still unclear at this point. The burst of Cer3 expression at 8-cell to gastrulation stages coincides with the time window when *C. elegans* embryo blastomeres are pluripotent[50, 51]. It is likely that expression of LTR-derived transcripts in early embryonic founder cells are functional and regulated event. In the context of development, early embryonic transcriptional relaxation is following by germline transcriptional relaxation. It is possible that the two burst of transcriptional relaxation are inter-dependent because of the unbroken physical link between germline and embryos.

An outstanding question in the field of nuclear RNAi is how epigenetic silencing signals are transmitted to the next generation via the germline? We found that Cer3 transcripts are absent in the gametes, therefore cannot be the physical carriers of the transgenerational epigenetic memory. Instead, Cer3 is transiently transcribed in early embryos. We recently characterized that the establishment of nuclear RNAi silencing at both endogenous targets and exogenous dsRNA targets is a transgenerational process (Kalinava unpublished data) [35], highlighting the importance of germline to embryo transmission in potentiating strong nuclear RNAi silencing. Another related study reported that early exposure to exogenous dsRNA during embryogenesis triggers strong nuclear RNAi silencing in adult pharyngeal muscle at the same generation, suggesting that the early embryo stage is a critical time period of establishing nuclear RNAi silencing [52]. It will be of interest to further investigate if the short activation of Cer3 at *C. elegans* early embryogenesis stages is required to re-establish Cer3 small RNA biogenesis and epigenetic silencing, which may also provide immunities against similar foreign genetic elements at each generation.

### Cer3 as a model system to study the complex regulation and function of heterochromatin

We previously found that Cer3 chromatin contains a high level of the repressive histone modification H3K9me3. The H3K9me3 at Cer3 is deposited through both HRDE-1-dependent and HRDE-1 independent mechanism. Intriguingly, the high level of H3K9me3 is completely dispensable for Cer3 repression in the wild type *hrde-1* background, indicating that the HRDE-1-dependent Cer3 repression can occur through a H3K9me3-independent mechanism. On the other hand, mutant animals with combined losses of H3K9me3 and *hrde-1* exhibit a much higher Cer3 expression than the *hrde-1* single mutant. Therefore, H3K9me3, likely the HRDE-1-independent one, functions synergistically with HRDE-1 to repress Cer3 transcription. Similar to *hrde-1* mutant animals, the enhanced Cer3 expression in the germline of *H3K9 hmt;hrde-1* compound mutants is largely restricted to the distal region of the adult gonads and the PGCs. Cer3 repression in the proximal regions of adult gonads *(e.g*., diplotene and diakinesis) is mediated by a mechanism(s) that is independent of HRDE-1 and H3K9me3.

In intestinal cells and early embryos, H3K9me2 histone methyltransferase MET-2 and nuclear RNAi factor HRDE-1 synthetically suppress Cer3 transcription. In *met-2 hrde-1*, the transient transcription of Cer3 in early embryos increased >7-fold, followed by Cer3 mis-expressed in intestinal cells in late embryos that persists to adult stage. In contrast, H3K9me3 histone methyltransferase SET-25 and putative histone methyltrasferate SET-32 are not required in Cer3 chromatin silencing in somatic lineage, highlight the differential requirements of chromatin-mediated Cer3 silencing in germline and soma. MET-2 belongs to SynMuv B heterochromain protein family. Many synMuvB mutants show enhanced RNAi (eri) and misexpression of germline genes in intestine and hypodermis[53–55]. We hypothesize that in *met-2*, loss of H3K9me2 in early embryos activates gene expression program characteristic of less differentiated germ cells (for example, PGCs). Both germline nuclear RNAi pathway and Cer3 LTR-retrotransopon gained transcription potential in *met-2* intestinal cells, but Cer3 is repressed by the co-expression of germline nuclear RNAi pathway. In *met-2 hrde-1* double mutant, defective germline nuclear RNAi pathway can no longer repress Cer3 in intestine and leads to Cer3 transcription activation in intestinal cells.

## Materials and Methods

### Worm strains

*C. elegans* strain N2 was used as the standard wild-type strain. Alleles used in this study were *hrde-1(tm1200), nrde-2(gg091), met-2(n4256), set-25(n5021), set-32(red11), gld-1(q485), glp-1(ar202), ruIs32 [pie-1::GFP::H2B + unc-119(+)], bcIs39 [lim-7p::ced-1::GFP + lin-15(+)], red3 [sf-hrde-1]*. JU533 is a *C. elegans* wild isolate (Finistère, France) acquired from Caenorhabditis Genetics Center (CGC). JU533 has Cer3/gag deletion that was confirmed by Sanger sequencing. Animals were cultured on NGM plates with *E. coli* OP50 as the food source in a temperature controlled incubator.

### Single Molecule Fluorescent *in situ* Hybridization (smFISH)

smFISH fluorescent probe sets were designed using Stellaris^®^ Probe Designer (Biosearch). Sequence information of each probe sets are in (Supplemental Material S1). The Cer3/gag smFISH probes were designed using a 2242-nt protein-coding sequence in the Cer3 *gag* gene. Cer8 has two full-length copies in *C. elegans* genome, with one of the copy having a Tc1 DNA transposon insertion in its internal sequence. The two copies are both endogenous nuclear RNAi targets and have non-homologous sequence around the Tc1 insertion site. We designed Cer8 smFISH probes using 2001-nt non-homologous sequence specific to Cer8 (C03A7.2), the copy that lacks Tc1 insertion. The different amount of Cer3/gag or Cer8 smFISH signals in wild-type and *hrde-1* mutant correlates with our previous RNA-seq and Pol II-ChIP analysis(refs), suggesting the probes are specific.

The embryo fixation protocol is adapted from Tintori et al[56]. Embryos were isolated from gravid adult worms by bleaching, resuspended in methanol (−20 °C), freeze cracked in liquid nitrogen, and then stored at −20 °C overnight. Larval fixation protocol is adapted from Ji et al. [57]. Developmental synchronized worms of desired stage were resuspended in fixation solution (4% paraformaldehyde in 1x PBS) and rotated at 19°C for 45 minutes. After fixation, the hybridization, washing, and mounting for both embryos and larvae were carried out using protocol described in Ji et al. [57]. Fixed samples were washed in wash buffer (10% formamide in 2xSSC) for 5 mins, and then incubated in 100ul of 125nM smFISH probe set in hybridization buffer (0.1g/ml dextran sulfate, 1mg/ml *Escherichia coli* tRNA, 2mM vanadyl ribonucleoside complex, 0.2mg/ml RNase free BSA, 10% formamid) at 30°C overnight. After hybridization, samples were washed in wash buffer at 30°C for 30 minutes, incubated in 50ng/mL DAPI in wash buffer at 30°C for 30 minutes, washed once in 2xSSC for 2 minutes at room temperature, and then stored and mounted in SlowFade (ThermoFisher).

Single color smFISH images were acquired using a DeltaVision Image Restoration Microscope system (Applied Precision Instrument) using a 100×/1.35 UplanApo objective and a Cool Snap HQ2 camera. Images were deconvoluted with the HuygenEssential software (Science Volume Imaging). Two color smFISH images were acquired using Zeiss Axiover200M Microsope using a 63x/ 1.3 objective and a cooled CCD camera (Q-imaging). Images were captured by MetaMorph software and then were deconvoluted by AutoQuant X3 software. Image J (NIH, http://imagej.nih.gov/ij/) was used for viewing and processing deconvoluted image data. ImageJ maximum projection was used to project z-stack images to a single plane. The fluorescent intensity of smFISH dots were >2-fold above background as expected [57]. smFISH dots in embryos were counted using smFISH quantification software StarSearch (Arujun Raj, University of Pennsylvania).

### CRISPR/Cas9-mediated genome editing

N-terminus SF tag (strep II-Flag) was inserted after the start codon of hrde-1 (c16c10.3) using CRISPR-Cas9-based genome editing. SF-HRDE-1 is anti-FLAG western blot positive. It expresses in germline nuclei as expected shown in anti-FLAG immuostaining(Sup Fig1A). CRISPR/Cas9-mediated genome editing was performed using protocol described in Arribere, et al. [58]

### Preparation of worm grinds

Synchronized young adult worms were prepared using the hypochlorite bleaching method described in[59]. Before grinding, synchronized young adult worms were harvested by washing off the plates with M9 buffer. Bacteria washed off along with the worms were separated and removed by loaded harvested sample to the top of 10ml 10% sucrose in a 15ml conical tube and centrifugation in a clinical centrifuge. The cleaned worm pellet at the bottom was immediately pulverized by grinding in liquid nitrogen with a mortar and pestle and were stored at −80 °C.

### Pull down SF-HRDE-1 bound endogenous siRNA

We used FLAG Immunoprecipitation kit (Sigma Catalog# FLAGIPT1) and followed the the manufacture’s protocol to pull-down SF-HRDE-1 and its bound siRNA. Briefly, the frozen worm grind was first lysed in 1ml of Lysis Buffer, 10ul of 100xHALT protease inhibitor (ThermoFisher), and 10ul of RNaseOUT (ThermoFisher). After incubating at room temperature for 15 minutes, the lysis were sonicated three times, 8 minutes each time, at 4°C using a Bioruptor (Diagenode) at high setting and 30 second on/off cycle. Then the lysis were centrifuged at top speed for 4 minutes at 4°C. The supernatant was mixed with ANTI-FLAG M2 affinity gel (washed and collected from 40ul resin provided by kit) and was rotated at 4°C overnight. On the next day, the sample was centrifuged at 8,200xg for 30 seconds at 4°C and the supernatant was removed. The resin was washed three times with 0.5ml of 1x Wash Buffer at 4°C. SF-HRDE-1 was eluted from resin with 15ug of 3xFLAG Peptide in 100ul of 1x Elution Buffer by rotating 30 minutes at 4°C.

### Nuclear and cytoplasm purification

Nuclei and cytoplasm factions were obtained using Nuclei PURE Prep kit (Sigma-Aldrich NUC-201) according to manufacture’s instruction. All procedure was performed at 4°C. Briefly, worm grinds were re-suspended in 10ml Lysis Solution with 1mM DTT and 0.1% Triton X-100. Proper grinding was confirmed by DAPI staining the lysis and microscopic examination. The lysis was mixed with 18ml 1.8M Sucrose Cushion, loaded on top of 10ml 1.8M Sucrose Cushion Solution, and ultra-centrifuged at 30,000xg for 45 minutes at 4°C in a SW28 rotor. The clear sucrose cushion layer was collected as cytoplasm fraction and used for RNA extraction immediately. The pellet of purified nuclei was wash with Nuclei PURE Storage Buffer, and was used as nuclear fraction for RNA extraction immediately.

### High-throughput sequencing

RNA-seq: Total RNA was extracted from nuclear faction or cytoplasm faction using Trizol reagent (Life Technologies). Ribosomal RNA(rRNA) was depleted using RNaseH and PAGE-purified DNA oligos (mixture of 110 Oligos, 50-nt) that are antisiense to rRNA. Briefly, 1ug of rRNA antisense oligos was mixed with 1ug of total RNA. The sample was denatured at 95 °C for 2 minutes, and then cooled at −0.1 °C /sec to 22 °C. rRNA was digested by 10 units of Hybridase Thermostable RNase H (Epicentre) in 1x RNaseH Reaction Buffer at 45°C for 30 minutes. DNA oligos were removed by DNaseI (NEB) according to manufacture’s instruction, followed by phenol: chloroform extraction. The yielded RNA was used to prepare RNA-seq library as described in [1]. We validated the subcellular fractionation by examining intronic reads, as nuclear RNA-seq libraries should contain more intronic reads than the cytoplasmic ones. This was indeed the case for large introns (2-10 kb), which accounted for 6.3% of all annotated introns (Fig S7).

#### sRNA-seq

Small RNA was enriched using the mirVana™ miRNA Isolation Kit (Life Technologies). Small RNA library was prepared from 1 μg small RNA by using a 5’-mono-phosphate-independent small RNA cloning procedure as described previously[1].

#### Multiplexing

Each library is barcoded with a unique 6-mer index located on the 3’ linker. Libraries of the same type (RNA-seq and sRNA-seq) and the same biological repeat were pooled together for HTS.

#### HTS instrument

Pooled libraries were sequenced on an Illumina HiSeq 2500 platform with the following specifications: rapid run mode, 50-nt single-end run, and index sequencing. De-multiplexed raw data in fastq format were provided by the sequencing service provider. Library information is listed in Supplemental Material S2.

## Acknowledgments

We thank B. Grant, A. Norris, H. Ushakov, E. Gavin, and S. Patel for technical assistance and discussions. Research reported in this publication was supported by the Busch Biomedical Grant and the National Institute of General Medical Sciences of the National Institutes of Health under award number R01GM111752. Some strains were provided by the CGC, which is funded by NIH Office of Research Infrastructure Programs (P40 OD010440). The content is solely the responsibility of the authors and does not necessarily represent the official views of the National Institutes of Health.

## Author Contributions

Conceived and designed the experiments: SG JN. Performed the experiments: JN SG NK SGM. Analyzed the data: JN SG. Wrote the paper: JN SG.

